# Orchestrating Single-Cell Analysis with Bioconductor

**DOI:** 10.1101/590562

**Authors:** Robert A. Amezquita, Vince J. Carey, Lindsay N. Carpp, Ludwig Geistlinger, Aaron T. L. Lun, Federico Marini, Kevin Rue-Albrecht, Davide Risso, Charlotte Soneson, Levi Waldron, Hervé Pagès, Mike Smith, Wolfgang Huber, Martin Morgan, Raphael Gottardo, Stephanie C. Hicks

## Abstract

Recent developments in experimental technologies such as single-cell RNA sequencing have enabled the profiling a high-dimensional number of genome-wide features in individual cells, inspiring the formation of large-scale data generation projects quantifying unprecedented levels of biological variation at the single-cell level. The data generated in such projects exhibits unique characteristics, including increased sparsity and scale, in terms of both the number of features and the number of samples. Due to these unique characteristics, specialized statistical methods are required along with fast and efficient software implementations in order to successfully derive biological insights. Bioconductor - an open-source, open-development software project based on the R programming language - has pioneered the analysis of such high-throughput, high-dimensional biological data, leveraging a rich history of software and methods development that has spanned the era of sequencing. Featuring state-of-the-art computational methods, standardized data infrastructure, and interactive data visualization tools that are all easily accessible as software packages, Bioconductor has made it possible for a diverse audience to analyze data derived from cutting-edge single-cell assays. Here, we present an overview of single-cell RNA sequencing analysis for prospective users and contributors, highlighting the contributions towards this effort made by Bioconductor.

## Introduction

Bioconductor [1] is a central repository for open-source software packages based in the R programming language [2] for the analysis and comprehension of high-throughput biological data. Since 2001, it has drawn together a rich community of developers and users from diverse scientific fields including genomics, proteomics, metabolomics, flow cytometry, quantitative imaging, chemoinformatics, and other high-throughput data.

Bioconductor supports the analysis of traditional bulk DNA, RNA, and epigenomic profiling assays [3–10] (Figure 1). Such bulk profiling technologies have yielded a wealth of important scientific insights (reviewed in e.g. [11–13]. However, a number of critical biological questions can be only answered at the single-cell level. Characterizing the extent of genetic heterogeneity within a tumor, identifying and characterizing rare cell populations with differential features, and defining the mechanisms of cell lineage differentiation, are all examples of biological questions that are intractable without single-cell approaches. Furthermore, revisiting questions that were previously tackled with bulk approaches can potentially provide new perspectives. Biotechnology is an ever-evolving field, with an ever-changing vocabulary (see **Box 1** for our definitions used throughout), and new single-cell experimental protocols and technologies such as single-cell RNA-sequencing (scRNA-seq) have emerged that can help tackle these unresolved questions.

**Figure 1:**
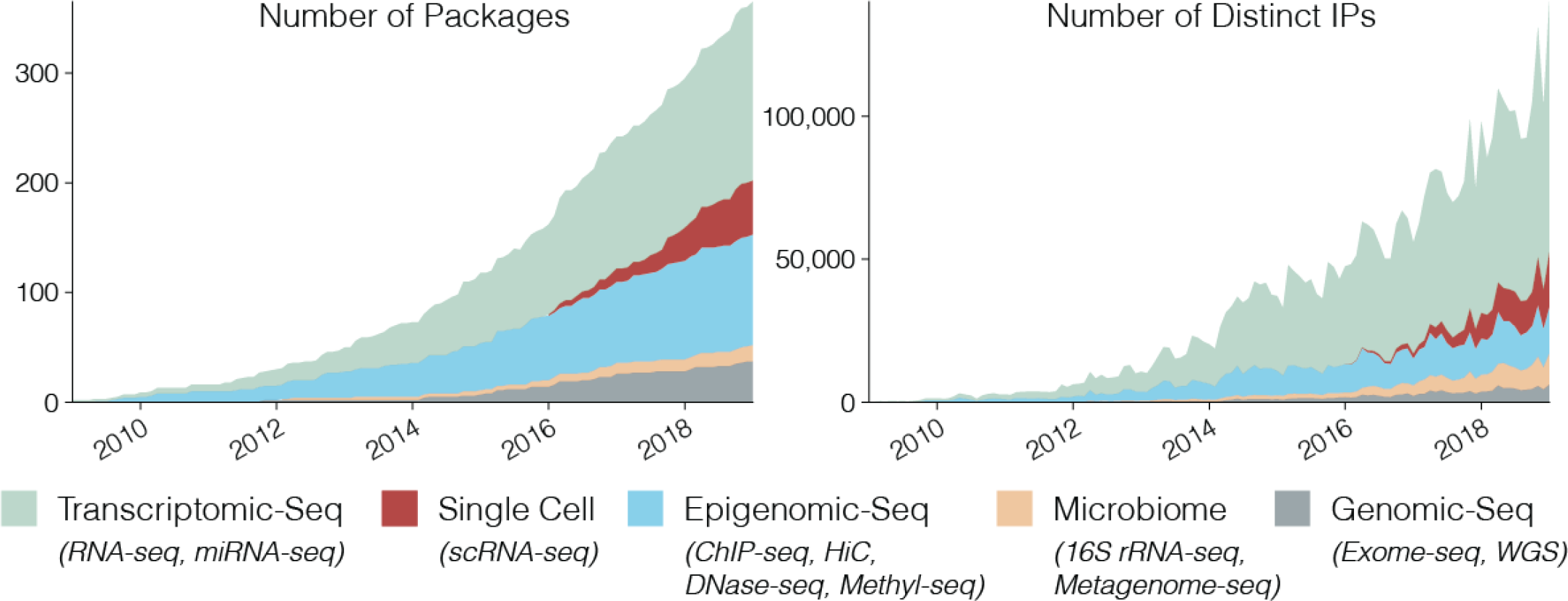
10 years of Bioconductor in the high-throughput sequencing era. Bioconductor software packages associated with the analysis of sequencing technology were tracked by the total number of packages (left) and the number of distinct IPs (data recorded monthly) visiting their online documentation (right) over the course of ten years. Software packages were uniquely defined by their primary sequencing technology association, with examples of specific terms used for annotation below in parentheses.

In addition, Bioconductor has been a pioneer for analyzing high-throughput data from single-cell cytometry assays that can be used to obtain samples from hundreds of thousands to millions of cells [14,15], but are comparatively low-dimensional, capturing around 20 to 50 features per cell. In contrast, the more recently developed single-cell genomics technologies are both high-throughput and high-dimensional, capturing thousands of traits within a single cell in an unbiased manner. To successfully derive biological insights from such work, two unique characteristics of the resulting data sets must be successfully dealt with. One is the increased scale on which samples, or cells, are measured, i.e. thousands to millions of cells within each data set, for example in the compendiums from the Human Cell Atlas [16]. In contrast, data derived from bulk assays, such as RNA-sequencing (RNA-seq) and chromatin immunoprecipitation sequencing (ChIP-seq), typically have samples on the order of tens to hundreds. A second distinctive feature of single-cell assay data is their increased sparsity, due to biological fluctuations in the measured traits or limited sensitivity in quantifying small numbers of molecules [17–19]. In addition, data derived from single-cell assays have revealed more heterogeneity than previously seen [20–27]. This has led to the rapid development of statistical methods to address the increased sparsity and heterogeneity seen in this data [28–31]. The profound increase in the complexity of data measured at the single-cell level, along with the continued increases in the number of samples measured, have precipitated the need for fundamental changes in data access, management, and infrastructure to make data analyses scalable to empower scientific progress. Specifically, specialized statistical methods along with fast and memory-efficient software implementation are required to reap the full scientific potential of high-throughput, high-dimensional data from single-cell assays.

Bioconductor has developed state-of-the-art and widely used software packages (Table S1) for the analysis of high-dimensional bulk assays, such as RNA-sequencing (RNA-seq) and high-throughput, low-dimensional single-cell assays, such as flow cytometry and mass cytometry (CyTOF) data. Therefore, Bioconductor is a natural home for software development for data derived from high-throughput, high-dimensional single-cell technologies. In particular, Bioconductor has developed state-of-the-art software packages (Table S1) and workflows (Table 1) to analyze data from such technologies (Figure 2). To help users get started with Bioconductor, we describe some of these packages and present a series of workflows (**Box 2**) to demonstrate how to leverage the robust and scalable Bioconductor framework to analyze data derived from single-cell assays. We primarily focus on the analysis of scRNA-seq data, although many of the packages mentioned herein are also generalizable to other types of single-cell assays.

Broadly, these new Bioconductor contributions provide profound changes in how users access, store, and analyze data, including: (1) memory-efficient data import and representation, (2) common data containers for storing data from single-cell assays for interoperability between packages, (3) fast and robust methods for transforming raw single-cell data into processed data ready for downstream analyses, (4) interactive data visualization, and (5) downstream analyses, annotation and biological interpretation. As a companion to this manuscript, we also provide an online book that provides extensive resources for running R and Bioconductor, and furthermore demonstrates select workflows corresponding to topics covered in this manuscript. The book can be found at: https://osca.bioconductor.org

#### Box 2. Key Definitions

***Sample***: a single biological unit that is assayed.

***Feature***: a trait of a sample that is measured. Examples include mRNAs in RNA-seq experiments, genomic loci for ChIP-seq experiments, and cell markers in flow/CyTOF experiments.

***Experiment***: a procedure where a set of features are measured for each sample; in this usage, typically involves multiple samples, possibly with varying conditions (e.g. treatments, time points).

***High-throughput assay***: an assay that captures and measures features from many samples. Examples include flow cytometry, CyTOF, and certain scRNA-seq platforms, which can quantify tens or hundreds of thousands to millions of cells. For this reason, in our review, most bulk assays are not considered high-throughput as they profile a limited number of samples.

***High-dimensional assay***: an assay that captures thousands or tens of thousands of features per single sample unit. In our review, high throughput assays such as flow cytometry are not considered high-dimensional as they profile a limited number of proteins.

***Bulk assay***: an assay that measures pools of cells to produce a set of measured features as a single observation unit per pool.

***Single-cell assay***: a technology where a single sample corresponds to a single cell; includes flow cytometry, CyTOF, and single-cell RNA-seq (scRNA-seq) across various platform technologies (plate-based, droplet, etc.).

#### Box 2. Getting Started

To accompany this manuscript, we have written a book that is freely accessible online and discusses in detail how to get started with using R and Bioconductor. The book covers installation, learning how to get help, and workflows covering specific case studies to illustrate the usage of R and Bioconductor based workflows. See the book at: https://osca.bioconductor.org

### Preprocessing sequencing data

The analysis of sequencing-based assays often begins with quantification of measured traits from raw sequencing reads. For high-throughput scRNA-seq assays, sequenced reads are typically aligned to the transcriptome and quantified to create a matrix of expression values for each cell across the features of interest (i.e. genes or transcripts) for further analysis. While much of the specific choices defining the workflow that generates processed data from raw data are often technology- or platform-dependent, we will briefly touch on the topic of alignment. A pipeline to process scRNA-seq data based on the R/Bioconductor software project is now available through the *scPipe* package [32], which uses the *Rsubread* Bioconductor package [33,34] to provide alignments within R. For droplet-based scRNA-seq technologies, such as 10X Genomics [35], the *DropletUtils* Bioconductor package can read in a matrix of UMI counts, which were produced from, for example, the *Cell Ranger* [35] 10X Genomics pipeline, where each row of the matrix corresponds to a gene, and each column corresponds to a cell barcode.

Outside of R, recent developments in alignment methods have produced a class of pseudoaligners capable of running on personal computers, such as the Salmon [36] and Kallisto [37] utilities. The *tximport* [38] Bioconductor package imports the results from these pseudoaligners as matrices into an R session. In combination with the *tximeta* [39] package, an instance of a well-supported Bioconductor class can be created. Common Bioconductor methods and classes are the foundation of the single-cell R/Bioconductor analysis software discussed herein. The specific infrastructure used to represent single-cell data is discussed in more detail in the next section.

### Data Infrastructure

A key focus and advantage of embracing Bioconductor workflows is the use of common data infrastructures. First and foremost, the implementation of the data containers is done with the aim of enabling modularity between packages and interoperability across packages. Hence, the containers are designed to support diverse workflows, while still being accessible to end-users. To this end, Bioconductor makes use of the S4 object-oriented programming style that allows for the creation of classes that standardize how data is stored and accessed. Furthermore, a single object or data container can contain a rich set of metadata annotation, as well as various forms of primary data essential for description, analysis, and portability. Bioconductor has recently focused its efforts on the creation of the *SingleCellExperiment* class to support single-cell data platforms, which is described in depth below.

### The *SingleCellExperiment* container

The *SingleCellExperiment* class is a lightweight data container for storing data from single-cell assays or experiments [40] that is sufficiently flexible to work with a diverse set of packages. Because the *SingleCellExperiment* class was developed as an extension of the *SummarizedExperiment* [41] class, the *SingleCellExperiment* class contains all the advantages, structure, and specially engineered features to accommodate large-scale data, with a number of additions that provide convenient methods and structures that are common to many single-cell analyses. Specifically, the inheritance of *SingleCellExperiment* class from the *SummarizedExperiment* class [41] has enabled the immediate use of previously developed methods, many of which are discussed by Huber *et al.* 2015 in a previous review of Bioconductor tools [1]).

**Figure 2:**
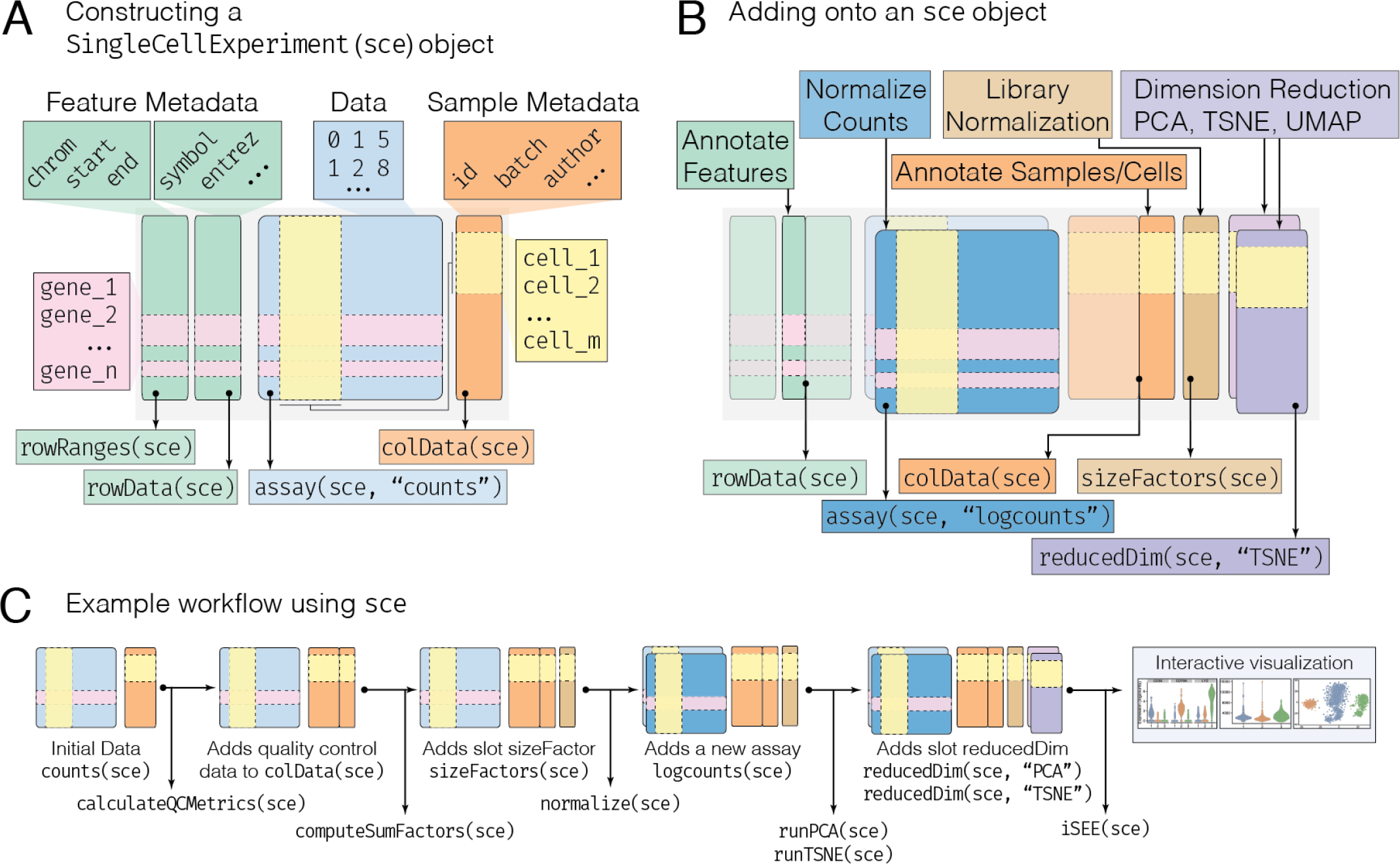
Overview of the *SingleCellExperiment* class and workflow. The *SingleCellExperiment* (sce) class and data container from the *SingleCellExperiment* package [40] stores multifaceted large-scale data and enables interoperability across Bioconductor packages. References to R functions and objects are shown in monospace (code) font. **(A)** A minimal sce object is constructed by supplying data such as a matrix of counts per cell as an assay (blue box), consisting of features, such as genes (rows), and cells (columns). Metadata describing the cells may also be supplied, wherein the cells are represented as rows and known characteristics of the cells are columns (orange box). Similarly, metadata describing the features may be added as well (green box). Each of these different types of data are stored in distinct parts of the sce object, which are referred to as slots. The data within each slot may be accessed programmatically via accessors named after their respective slot (arrows), such that rowRanges refers to feature metadata, colData refers to sample metadata, and assay refers to data. (**B**) Analysis using *SingleCellExperiment* (sce) compatible workflows appends data to the initial sce object. For example, calculating library normalization factors per cell creates a new slot (pink box). These can then used to derive a normalized count matrix, which is stored in the same assay slot alongside the initial counts data (dark blue box). The assay slot is thus capable of storing any number of data transformations. Cell quality metrics, which describe cell characteristics, are appended to the sample metadata slot colData. Finally, in a similar fashion to the assay slot, any number of dimensionally reduced representations of the data can be stored, residing in their own slot, reducedDim (purple box). **(C)** The sce object evolves throughout the course of a typical analysis, storing various metrics and representations derived from the initial data. For more information on the *SingleCellExperiment* class, see the *SingleCellExperiment* vignette (https://bioconductor.org/packages/SingleCellExperiment).

Made up of multiple compartments of information called *slots*, the *SingleCellExperiment* object holds various data representations (Figure 3). Primary data, such as count matrices representing sequencing read or unique molecular identifier (UMI) counts, are stored in the assays slot as one or more matrices (including sparse or dense Matrix [42] objects), where rows represent features (e.g. genes, transcripts, genomic regions) and columns represent samples, or in the case of single-cell experiments, cells. Furthermore, each row and column can be annotated with a rich set of metadata. For example, row metadata is stored in the rowData slot and could include Entrez Gene IDs [43] and GC content. Further, for rows corresponding to genomic features, a special rowRanges slot can be created to hold genome coordinates. Column metadata is stored in the colData slot and can contain information about sample-level characteristics. Additional column metadata can be added to this slot as well - for example, summary quality control statistics such as the total number of reads per cell. This column metadata is particularly useful for subsetting the data, such as removing cells with a low number of reads. Separately, the sizeFactors slot also refers to columns (cells) and contains information necessary for normalization. A recent innovation that is specifically designed for single-cell data is the *reducedDims* slot that contains low-dimensional representations of data, such as principal components analysis (PCA), *t*-Distributed Stochastic Neighbor Embedding (*t*-SNE) [44], or Uniform Manifold Approximation and Projection (UMAP) [45].

**Figure 3:**
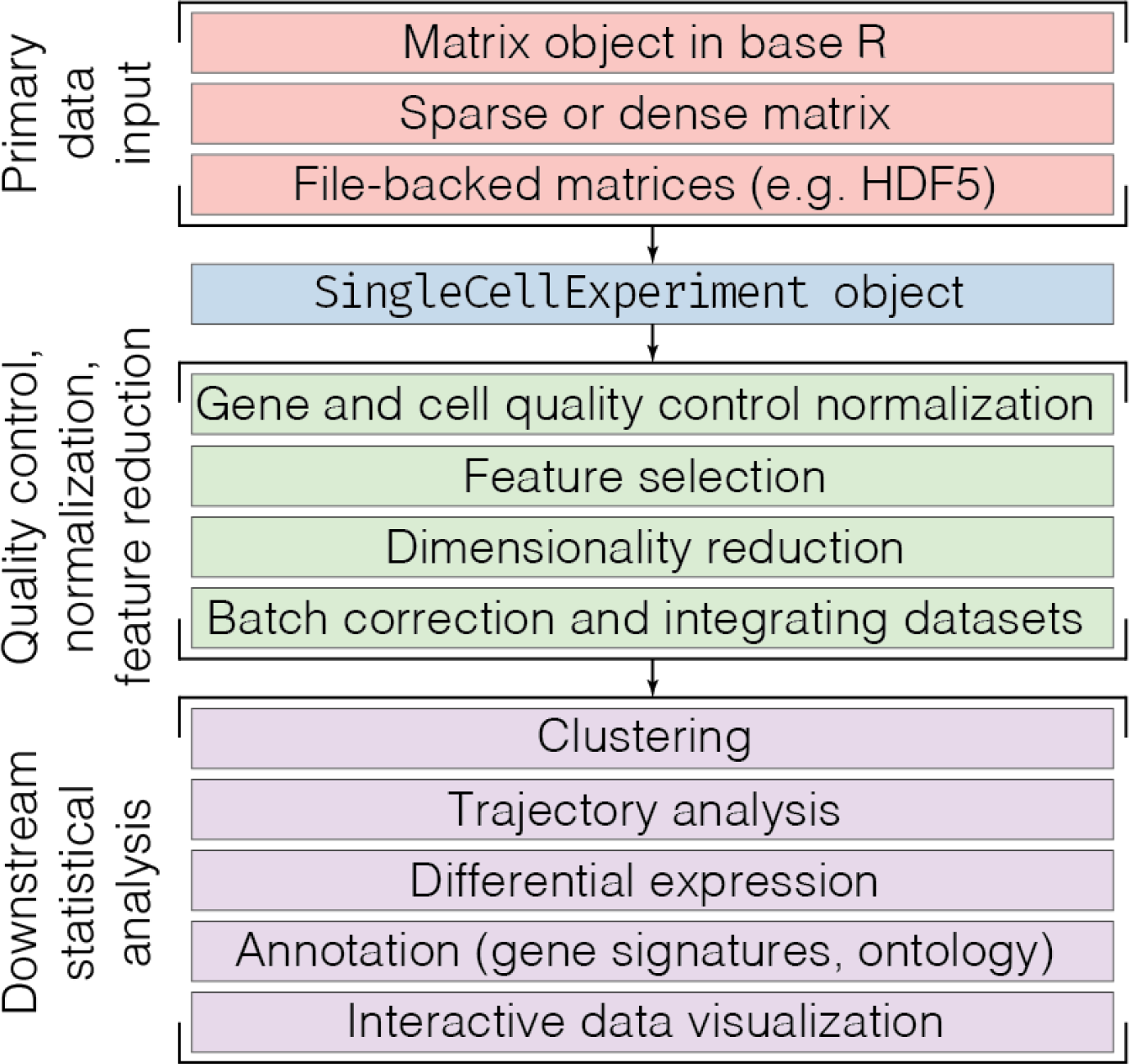
Bioconductor workflow for analyzing single-cell data. Primary data is collected and coalesced into a matrix representation, which can exist in various forms (top, red box). This data is then used to create a *SingleCellExperiment* object, which contains both primary data and metadata essential to understanding the experimental setup (blue box). The *SingleCellExperiment* object is then transformed through preprocessing workflows that ultimately produce a cleaned expression matrix (green box). This result can then be used in various downstream statistical analyses (purple box, bottom). This paper is structured to roughly follow this analytical workflow from top to bottom.

Lastly, throughout the various slots of the *SingleCellExperiment* class, disk-backed representations of the data (e.g. HDF5) are supported to enable analyses that would otherwise be impossible to perform due to memory constraints. Recent software innovations - both in data handling and processing - are required to make full use of this capability, and are demonstrated in our online supplement (Table 1).

Altogether, these different types of primary data and metadata slots reside in a single data container, allowing for a portable, full representation and annotation of the data alongside continual validity checks to prevent malformed data input. This has made the *SingleCellExperiment* data container the foundation of many packages oriented toward single-cell analysis available today in Bioconductor, providing seamless interoperability and facilitating the development and usage of cutting-edge computational methods, from initialization of data containers to the end stages of analysis.

In addition, recent advances in technology and protocols allow the simultaneous measurement of genetic, epigenetic, and transcriptomic information from the same cells [46–52]. The *MultiAssayExperiment* [53] package integrates heterogeneous data types that may be individually represented by *SingleCellExperiment*, *DelayedArray*, or other standard R/Bioconductor data structures.

### Quality Control and Normalization

After quantifying a measured trait from a single-cell assay and creating a *SingleCellExperiment* object, one of the first steps in analyzing the data is to identify, remove, and correct for low quality features or samples. In the analysis of scRNA-seq data, this typically translates to transforming a raw count matrix to a processed expression matrix ready for downstream analyses, such as clustering or differential expression. This process includes filtering out low-quality cells, selecting for informative features, applying cell and gene-specific normalization methods to remove cell and gene-specific biases, and adjusting for known covariates as well as latent sources of variation. In this section, we discuss these steps in greater detail, as well as methods for integrating data from multiple single-cell experiments.

### Cell and Gene Quality Control

For droplet-based scRNA-seq technologies such as 10X Genomics [35], the *DropletUtils* package can be used to perform key quality control tasks such as distinguishing empty droplets from cells [54] and reducing the effects of barcode swapping [55]. The *scater* [56] package automates the calculation of a number of key quality control metrics. Amongst these, the library size (the total number of read or UMI counts across all genes in a given cell), the proportion of counts assigned to spike-ins [57] or mitocondrial genes, and the number of genes detected are commonly used to remove low quality cells. Furthermore, genes can be annotated with their abundance measurements, such as average expression, or frequency of detection, enabling the removal of uninformative genes [58]. The *simpleSingleCell* [59] Bioconductor workflow demonstrates how to use these packages to apply such quality control metrics for both read counts and UMI counts scRNA-seq data.

### Gene Expression Normalization

Though many normalization procedures exist for data derived from traditional bulk assays [3, 7, 9, 60–62], using data derived from single-cell assays, such as scRNA-seq, introduces new challenges that require cell and gene-specific biases to be eliminated prior to downstream analyses that depend on explicit gene expression value comparisons, such as differential expression analyses [19]. These biases can be reduced through the use of cell- (and possibly gene-) specific scaling factors, also known as size factors, which are used to make cells with different properties comparable. The family of approaches discussed herein explicitly address technical, experimental, and biological factors to make cells within a single scRNA-seq experiment comparable through the calculation of a corrected or “clean” gene expression matrix.

The *SCnorm* [63], *scran* [29] and *scater* [56] packages provide standalone normalization methods whose results can then be used in any downstream analysis. The *SCnorm* package estimates and removes genespecific variability due to sequencing read depth, along with accounting for other feature-level biases such as GC content and gene-length effects. The *scran* package calculates cell-specific size factors used for normalization in a manner that accounts for population heterogeneity by first clustering groups of cells with similar expression profiles. Finally, *scater* can calculate size factors from the library sizes. However, these approaches can only account for intrinsic factors that can be deduced directly from the data, and thus cannot correct for experimental factors such as batch effects.

Alternatively, there are approaches that propose statistical models to address not only normalization, but also other analyses such as dimensionality redution or differential expression. The Bioconductor packages *BASiCS* [64, 65], *zinbwave* [31], and *MAST* [28] are not specific for normalization, but each provide unique statistical frameworks tailored for scRNA-seq data. Such frameworks have the capacity to adjust not only for intrinsic technical factors, but also for known artifacts such as time, treatment, and batch effects. Furthermore, these technical or experimental factors are typically unwanted variation, however at times, they may be potentially confounded with interesting biological factors or variation [19].

To compare the effects of different normalization strategies, the *scone* package [66] can be used to explore the results of various strategies and parameterizations. To learn more about scRNA-seq normalization methods, including a thorough discussion on the use of synthetic control genes such as ERCC spike-ins [57], we refer the interested reader to reviews by McDavid et al. [67] and Vallejos et al. [68].

### Adjusting for Cell-Cycle Heterogeneity

Unless a cell cycle synchronization step is performed, the individual cells in a population will be at different stages of the cell cycle when the traits of interest are analyzed. Thus, a special type of normalization that is unique to single-cell data is adjusting for the biological effect of cell cycle phase on gene expression profiles [69]. The biological heterogeneity induced from differences in cell-cycle phases across cells is often considered a source of unwanted, technical variation, as they can obscure primary biological effects or conversely be highly correlated with biological effects [70]. For this reason, computational methods may be needed to deconvolute cell-cycle heterogeneity. For instance, latent variable models have been proposed to remove cell-cycle variability [70]. Within the Bioconductor framework, the *scran* [59] package implements a trained cell-cycle classifier using known cell cycle genes in humans and in mice to assign G1, G2/M, and S scores to cells. These scores can then be used as covariates to regress out the effect of cell cycle. In addition, the *Oscope* [71] package can be used to identify oscillators, or oscillating genes, when a single cell’s mRNA expression is oscillating through cell states.

### Integrating Datasets

As scRNA-seq approaches continue to gain in popularity and decrease in cost, large-scale projects that combine and integrate independently generated datasets will become standard [72]. However, overcoming the inherent batch effects [73] of such approaches presents a unique challenge. While linear or generalized linear modeling frameworks can be used to integrate disparate datasets, the performance of these frameworks in the scRNA-seq context may be sub-optimal, due to their underlying assumption that the composition of cell populations is either known or identical across groups of cells [74]. While this assumption can be addressed through explicit consideration of similar clusters of cells, new approaches to integrate datasets from distinct single-cell experiments that are largely independent of linear models, and which do not even rely on producing a corrected gene expression matrix, have been developed to address this issue.

One approach involves the identification of the most biologically similar cells between batches using mutual nearest neighbors (MNNs) [74], which is implemented in the *scran* [59] and *batchelor* [75] packages. The MNN-corrected data can be directly used in downstream analyses, similar to other dimensionality reduction approaches, such as clustering or trajectory analyses. The *scMerge* package draws upon a similar approach, but instead uses mutual nearest clusters to merge datasets [76]. However, *scMerge* requires a user to supply the number of clusters *a priori* or have annotated cell types to run in unsupervised or semi-supervised mode, respectively. Similar to MNN in *scran* and *scMerge*, the *scmap* package [77] performs an approximate nearest neighbor search to project cells from a query dataset or experiment onto the cell types (clusters) or individual cells in a different dataset or experiment. Alternative implementations based on unsupervised deep learning methods such as the *scAlign* package [78] have also recently been proposed. Finally, several data integration methods developed for bulk assays are available in the *mixOmics* [79] package and have shown good performance on single-cell data [80].

Overall, the approaches that have been tailored for scRNA-seq and explicitly address dataset integration have performed well with respect to ameliorating batch effects in recent benchmarks comparing against traditional statistical modeling frameworks [80]. However, care should be taken in utilizing corrected gene expression matrices, and further validation should be applied to ensure the result is sufficiently free from artifacts. The integration result is especially useful for clustering and visualization. Statistical frameworks can thus then be applied on the (raw) counts data in service of downstream methods such as differential expression, which become more reliable thanks to the integration approach separating different cell types.

Beyond simply addressing batch effects, one particularly exciting application of these integration approaches is in facilitating comparisons of *de novo* scRNA-seq data with published reference compendiums such as the Human Cell Atlas [16] or, for the model organism Mus musculus, *Tabula Muris* [81, 82]. Such approaches may not only facilitate cross-comparisons, but furthermore enable the annotation of *de novo* datasets on the basis of previously annotated references (see the *Annotation* section for a broader discussion).

In our online supplement, we demonstrate an example of combining datasets derived from different sources using the integration approaches described above (Table 1).

### Feature Selection

In most experiments, only a subset of genes drive heterogeneity across the population of cells profiled within a single-cell experiment. Feature selection is the process of identifying the most informative features or genes that drive the biological variation, such as genes with high variance from biological effects, rather than technical effects. In the analysis of scRNA-seq data, feature selection is an important step because it reduces the computational burden and removes noise in downstream analyses.

In cases with known groups of cells, correlation-based approaches [83] or the identification of differentially expressed genes [84] across groups can be used to select key features. However, in practice, these groups of similar cells are not often known *a priori* unless the experimental design contains sorted populations or other markers to designate known cell populations, such as engineered gene constructs. More commonly in scRNA-seq data, feature selection is either: expression-based, selecting genes with a high overall mean expression across cells; variance-based, selecting highly variable genes relative to overall mean expression [59, 64, 65]; dropout-based, selecting high dropout genes relative to overall mean expression [84]; deviance-based, selecting genes based on how well each gene fits a null model of constant expression across cells [85]; or a mixture of these strategies. For recent reviews comparing feature selection methods for scRNA-seq data, including the concordance across different approaches, see Yip *et al.* [86] and Andrews *et al.* [84].

### Dimensionality Reduction

Cutting-edge single-cell assays can potentially measure thousands of features genome-wide, in hundreds of thousands to millions of cells. With data on this scale, many types of analyses quickly become computationally intractable. While feature selection can mitigate this high-dimensional problem to a certain extent, it is often insufficient in reducing the complexity of single-cell data. Dimension reduction approaches elegantly resolve this dilemma by creating low dimensional representations that nonetheless preserve meaningful structure. The end result can then be subsequently used for data visualization as well as downstream analyses such as clustering and trajectory analysis.

The *SingleCellExperiment* container has a dedicated slot, reducedDims, for holding such reduced dimension representations of single-cell data. This slot is used by single-cell Bioconductor packages to provide uniform storage and access of linear and non-linear reduced representations of the data. For example, the *scater* [56] package uses this slot to store and visualize reduced dimension representations of single-cell data after applying dimensionality reduction methods. These include the top principal components (PCs) after performing principal components analysis (PCA), the *t*-Distributed Stochastic Neighbor Embedding (*t*-SNE) components [44] using the *Rtsne* [87] R package, the uniform manifold approximation and projection (UMAP) components [45] using the *umap* [88] R package, and diffusion maps [89] using the *destiny* Bioconductor package, respectively. In addition, the *BiocSingular* [90] package provides access to both exact and approximate singular value decomposition (SVD) methods for developers of Bioconductor packages to implement various forms of SVD within their own package. To improve the speed of computations in SVD methods, *BiocSingular* uses the *BiocParallel* [91] Bioconductor framework to parallelize operations.

The *zinbwave* [31] Bioconductor package takes an alternative approach, calculating a model-based dimensionality reduction based on the ZINB-WaVE model [31] tailored for zero-inflated count data that allows for adjustment for confounding factors, as described above. In a similar fashion to *scater*, the *zinbwave* [31] package works seamlessly with counts from *SingleCellExperiment* objects, and can store its results in the reducedDims slot for downstream analysis.

### Downstream Statistical Analysis

The choice of statistical analyses and workflows can differ greatly depending on the specific goals of the investigation. Following preprocessing of the data as described above, here we illustrate how the Bioconductor framework can be used to answer a variety of biological questions from single-cell data, using tools that are interoperable with the *SingleCellExperiment* class and scale with the number of cells. Also, we provide an online workbook (**Box 2**) that provides workflows and case studies for users on how to perform many of these downstream analyses with single-cell data.

#### Clustering

Data derived from single-cell assays have enabled researchers to unravel tissue heterogeneity at unprecedented levels of detail, enabling the identification of novel cell types, as well as rare cell populations that were previously unidentifiable using bulk assays [92–94]. Unsupervised clustering – the process of grouping cells based on a similarity metric without a known reference – is a fundamental step in deconvoluting heterogeneous single-cell data into clusters that relate to biological concepts, such as discrete cell types or cell states. Clustering is also essential for other analyses, such as differential expression, in order to identify distinct cellular sub-populations. While, the degree of the separation between the detected clusters is relevant to the robustness and confidence in the clusters, it is more important to think about using clustering methods to yield some hypothesis for experimental validation.

While many algorithms and software packages have previously been used to cluster data from bulk assays and single-cell flow cytometry [95], the complexities of scRNA-seq data pose unique challenges for clustering tasks, specifically a richer feature space, large numbers of cells, and data sparsity [96]. To address these challenges, Bioconductor has developed software packages that incorporate recent advance in nearest-neighbors and clustering algorithms that improve computational efficiency through approaches such as using approximate methods instead of exact methods, thereby trading an acceptable amount of accuracy for vastly improved runtimes. For example, the *BiocNeighbors* package [97–100] can be used to search for nearest neighbors and then a shared nearest neighbor graph using cells as nodes can be built using the *scran* [59]. Further, approximate methods have the advantage of smoothing over noise and sparsity, and thus potentially providing a better fit to the data [101]. Other approaches for improving computational speed include implementing specially designed versions of classical algorithms that support parallelization, enabling multicore and/or multinode processing [91].

Two implementations of unsupervised clustering frameworks come from two Bioconductor packages, *SC3* [102] and *clusterExperiment* [103], which calculate consensus clusters derived from multiple parametrizations. The *SIMLR* package [104] uses kernels to learn a distance metric between cells tailored to improve sensitivity in noisy single-cell data. For large data that require a file-backed representation such as a HDF5 file, the mini-batch *k*-means algorithm [105] has been implemented in the *mbkmeans* package [106], efficiently replicating the results of classical *k*-means clustering. For experiments with control genes present in the form of spike-in controls, the *BEARscc* [107] package estimates cluster variability due to technical noise [57]. For evaluating clustering performance across methods and parameter spaces, the results of these various methods can be assessed quantitatively and visually using the *SC3*, *clusterExperiment*, and *clustree* [108] packages. Together, these packages directly support the *SingleCellExperiment* class, allowing for seamless interoperability across a host of clustering methods.

Amongst these methods, the *SC3* package was found to be a top performer in two separate benchmarking studies [109, 110]. However, it should be noted that for the methods described above, only *SC3* was tested in the benchmarking studies. Thus, care should be taken in the choice of clustering method, as each method may excel in different contexts, and results should always be evaluated over various instantiations. For a demonstration of implementing clustering approaches, we refer the interested reader to our online supplement (Table 1).

#### Differential Expression

Given identifiable groups of cells, differential expression analysis can be used to identify features that uniquely distinguish each of the biological groups (clusters) of cells. The results from this analysis can then be used to identify the cell populations present. Another application involves comparing cells within a given population across various conditions, such as time or treatment. In both cases, it is important to correct for confounders such as systematic batch effects [19, 111]; furthermore, it is important to consider how these effects may drive clustering, and thus confound differential expression analyses from the start.

The distinct challenges described in the previous section for scRNA-seq data, namely data size and sparsity, have also spurred many methodological and computational developments in Bioconductor for identifying differentially expressed features between biological groups. Moreover, multiple recent benchmark papers have highlighted that many of the top performing scRNA-seq differential expression tools are Bioconductor packages (e.g. MAST and edgeR, mentioned below) [112–114].

Across these differential expression methods, two general approaches stand out. The first approach retrofits frameworks initially designed for bulk RNA-seq analysis, namely the *edgeR* [3, 62], *DESeq2* [7], or *limma* [115] frameworks. In this approach, the *zinbwave* [31] package can be used to model the single-cell data as arising from the zero-inflated negative binomial distribution (ZiNB) to account for the sparsity inherent to scRNA-seq data [116]. Namely, the *zinbwave* package downweights the excess zeros observed in scRNA-seq data in the dispersion estimation and model fitting steps, thereby enabling improved differential expression analysis. While this approach is slightly more complex at the onset, the *edgeR*, *DESeq2*, and *limma* packages have the benefit of being well-supported, robust, and with ample documentation.

The second class of approaches is uniquely tailored for single-cell data because the statistical methods proposed directly model the zero-inflation component, frequently observed in scRNA-seq data. These methods explicitly separate gene expression into two components: the discrete component, which describes the frequency of a binary component (zero versus non-zero expression), and the continuous component, where the level of gene expression is quantified. While all the methods mentioned herein can test for differences in the continuous component, only this second class of approaches can explicitly model the discrete component, and thus test for differences in the frequency of expression. To do this, the *MAST* [28] package uses a hurdle model framework, whereas the *scDD* [117], *BASiCS* [64, 65], and *SCDE* [18, 118] packages use Bayesian mixture and hierarchical models. Together, these methods are able to provide a broader suite of testing functionality and can be directly used on the scRNA-seq data contained within a *SingleCellExperiment* object.

For a guide in performing differential expression analyses, we refer the interested reader to our online supplement (Table 1).

#### Trajectory Analysis

In contrast to comparing distinct groups of cells within an experiment as described above, heterogeneity in some cases may be better explained as a continuous spectrum, arising due to processes such as cell differentiation. A specialized application of dimension reduction, trajectory analysis - also known as pseudotime inference - uses a broad array of phylogenetic methods to order cells (e.g. by beginning state, intermediate state, end state) along a trajectory of interest, such as a developmental process occurring over time (see [119, 120] for further discussion). From this inferred trajectory, it is possible to identify, for example, new subsets of cells, a differentiation process, or events responsible for bifurcations, such as branch-points, in a dynamic cellular process [121, 122]. As trajectory inference is intractable with bulk data, this type of analysis presents an exciting new avenue for methods development in single-cell applications.

New developments in trajectory inference methods have greatly expanded their capabilities. Whereas early versions required more guidance from the user - such as an expected topology - modern approaches have largely minimized the need for extensive parametrization. Furthermore, modern methods have led the development of novel statistical applications that can test for significant gene expression changes along a continuum and at branch points [123, 124].

In a recent review by Saelens et al. [125], they created a wrapper package - *dynverse* - to perform an extensive benchmarking of trajectory inference methods using both real and simulated datasets. From their evaluation, which included both performance aspects and qualitative features such as documentation, four of the five top performers were the Bioconductor packages *slingshot* [126], *TSCAN* [30], *monocle* [123, 124, 127], and *cellTree* [128]; the fifth was the *SCORPIUS* [129] package currently hosted on CRAN.

However, each of the methods above can produce drastically different results. This is largely due to the inherent complexity of the task and choices made by the authors of the packages in setting sensible default parameters, which may favor a specific type of topology. Therefore, it is essential to extensively test a suite of methods and parametrizations to assess the robustness of results from trajectory analysis methods. To facilitate such testing, the *monocle*, *slingshot*, and *CellTrails* (a newer method that as of this writing has not been tested by Saelens et al.) packages provide explicit support for *SingleCellExperiment* objects. For a more in-depth discussion on methods and benchmarking of trajectory analysis methods, we refer the interested reader to the previously mentioned paper [125] and a recent review by Tanay et al. [130].

### Annotation

One of the challenges of working with high-throughput data, such as that typically encountered in scRNA-seq, is the characterization of the data into familiar terms. This has led to the development of approaches that characterize expression changes of individual genes in the context of gene sets, often termed “gene signatures”. The basic structure of all gene signatures is that they provide a list of genes with a shared biological context. This context can be derived from myriad sources - from experimentally derived differential expression analyses to manually curated compilations of genes involved in metabolic and molecular pathways. More complex versions of gene signatures can also possess additional characteristics, such as quantitative or qualitative metrics, or even relationships between the elements of a given signature that produces a network representation. Quantifying the enrichment of these signatures for significant changes in expression has been a hallmark of functional annotation. In addition to this classical approach, novel methodologies developed in the era of scRNA-seq now bypass the usage of gene signatures, applying a data-centric approach to the annotation of *de novo* data using a reference dataset.

Here, we cover how such publicly available gene signatures can be applied to quantify the enrichment of gene signatures in scRNA-seq, methodologies that rely on reference data for annotation, and finally discuss a specialized application of these two approaches in assigning cell labels to single-cells or clusters of cells within an scRNA-seq experiment.

#### Accessing Public Gene Signatures

The characterization of gene signatures has made great strides through the coordinated efforts of many groups, which have pioneered strategies for standardizing cell type representations and developing statistical methods for the identification of necessary and sufficient markers for cell types and functional signatures [131,132]. The field has generated countless public knowledge-bases from which such signatures can be used for downstream enrichment analysis. These databases vary in their approach to the definition of signatures - some are based on experimental approaches, such as differential expression studies between conditions in the immunological module of MSigDB [133], whereas others rely on curated knowledge of well known molecular functions and biological processes, such as KEGG [134], Reactome [135], and Gene Ontology (GO) [136]. While covering the various available knowledge bases is outside of the scope of this manuscript, we encourage the use of programmatically accessible databases (e.g. those with application programming interfaces such as REST) and data packages which provide ready to use annotation data in order to facilitate reproducible analyses.

#### Gene Signature Enrichment

In this section, we focus on methods which rely on explicit gene signatures, either from manually curated or experimentally derived sources akin to those described above. Similar to identifying differentially expressed genes in single-cell data, there are two general approaches to test for an enrichment of genes in functional gene signatures. The first approach adapts existing gene set analysis methods originally developed for the analysis of microarray and bulk RNA-seq, such as *GSEA* [133] (or a fast implementation of pre-ranked GSEA in the *fgsea* package [137]), *goseq* [138], and *PADOG* [139], using observational weights to account for the excess zero observed in scRNA-seq data [116]. For a comprehensive set of available gene set analysis methods available on Bioconductor, see the *EnrichmentBrowser* package [140], which facilitates the usage of 10 different gene set based methods, with further options to combine resulting gene set rankings across methods. The second approach is a set of enrichment methods specifically tailored for scRNA-seq data. The *MAST* [28] package implements a competitive gene set test that accounts for inter-gene correlation using the hurdle model whereas the *AUCell* [141] package scores the activity level of gene sets using a rank-based scoring method and computes a gene set activation scores for each cell. Finally, the *slalom* [142] package uses a factorial single-cell latent variable model to explain variation in an scRNA-seq data set as a function pre-annotated gene sets.

#### Data-centric Enrichment Methods

A complementary approach to using published gene signatures relies on learning such signatures *de novo* from reference data. Such approaches, while still nascent, have the potential benefit of being able to characterize biological processes through the use of more comprehensive, quantitative signature definitions, as they go beyond just a list of genes. For example, the *scmap* [77] package projects cells from an scRNA-seq experiment onto the cell-types or individual cells identified in a different experiment. Similarly, the *scCoGAPS* [143,144] package generates gene expression signatures and then maps the learned signatures onto new datasets to learn shared biological characteristics.

#### Labeling of Cells

A specialized application of annotation methods in the analysis of scRNA-seq data is in automating the classification of unknown cells to known cell types. To accomplish this, a source of prior knowledge is first required. This can be in the form of a marker panel or a well-annotated reference scRNA-seq dataset, reflecting the division between gene signature and data-centric enrichment methods described above. Secondly, the choice of method will depend on the desired resolution of the annotation, as the labels are assigned either at the single-cell level or on (predefined) clusters of cells.

Methods that rely on reference scRNA-seq datasets that have been annotated *a priori* at the level of clusters, and thus apply data-centric enrichment approaches, possess some key advantages. Chiefly, by defining the labels at the cluster level from the reference dataset, e.g. from pools of cells, issues with sparsity can be overcome, and the level of uncertainty (variability) in the characteristic signature quantified. However, the definition of clusters, especially when it comes to defining biologically meaningful clusters, is an inherently empirical process, and thus any results relying on clusters - either from the reference dataset or the *de novo* dataset - will be subject to this initial bias. Ameliorating this, the definition of the characteristic signature pertaining to each reference cluster is much more quantitative than traditional manually curated gene signatures. Methods that adopt a cluster-centric labeling approach include the *celaref* [145] and *scmap* [77] packages. Interestingly, the *scmap* package also possesses the capability of annotating a *de novo* scRNA-seq experiment at the single-cell level (while still using the reference scRNA-seq clusters), thus removing one layer of potential bias arising from clustering.

Conversely, approaches that rely on manually curated panels have the benefit of being well-defined based on prior knowledge. Furthermore, they are easily adaptable, as the user can readily tweak the marker panel - extending or shortening signatures, as well as adding or removing cell type definitions. Naturally, such manual definitions also possess some disadvantages compared to data-centric definitions - for example, marker panels often rely on protein level characterization that may not be applicable to scRNA-seq, and these panels are usually much more limited in scope. Thus, it is important to apply such panels in a formalized manner, and one such package that takes this approach comes from *cellassign* [146], which labels individual cells from a *de novo* scRNA-seq experiment.

### Accessible and Reproducible Analysis

The richness of data from single-cell assays has immensely increased the space of possible data exploration. Oftentimes however, bespoke visualization frameworks to communicate results are limited in scope or may lack essential infrastructure that ensures reproducibility over time. To address these challenges, the Bioconductor community has embraced both software solutions and best practices, as described below.

**Figure 4:**
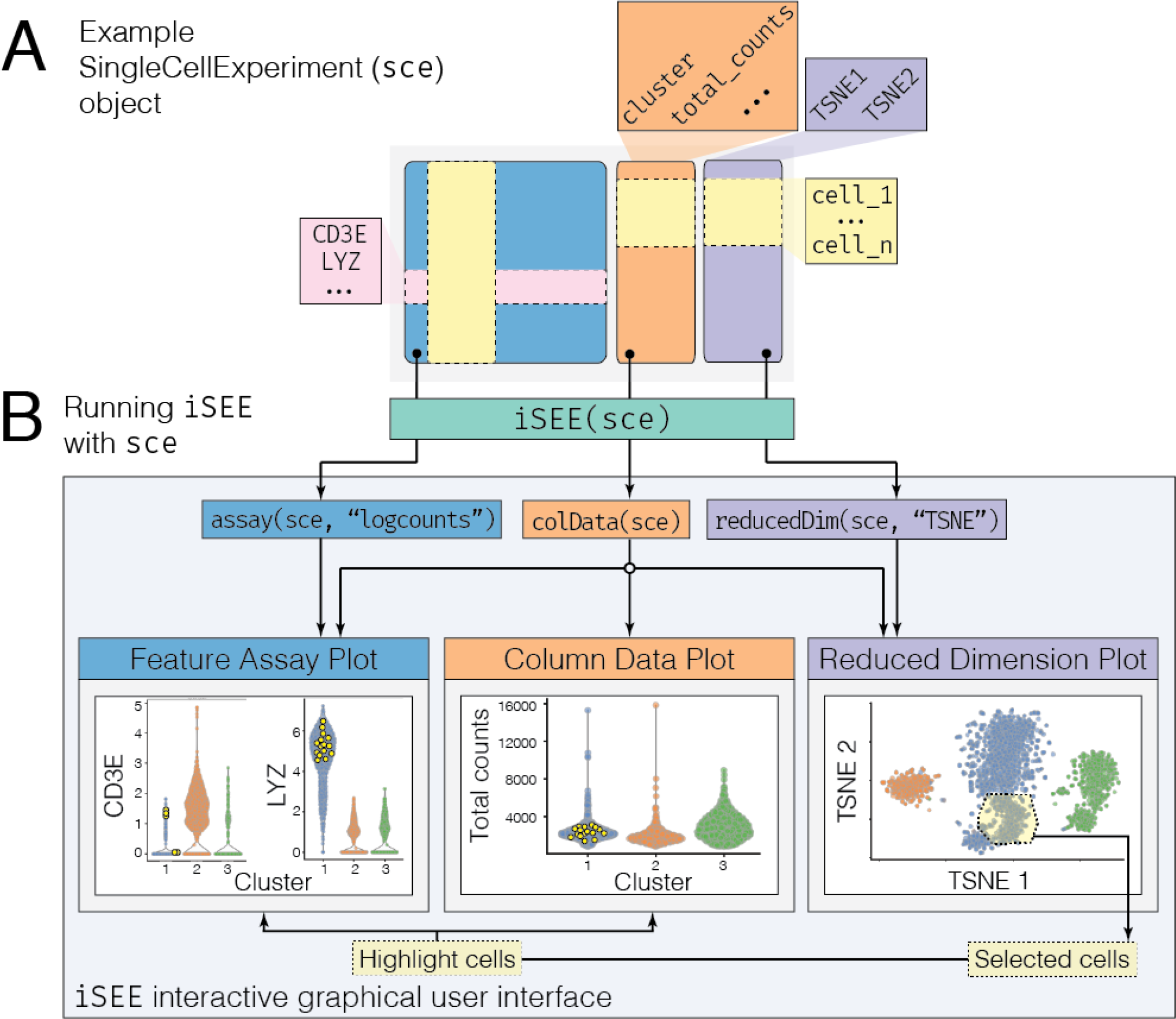
Interactive Data Exploration in Bioconductor. **A** An example *SingleCellExperiment* object, sce, is shown with a normalized expression assay that has been log-transformed (`logcounts`), sample metadata with a column representing the cluster label for each cell amongst other quality control metrics such as `total_counts`, and a reduced dimension `TSNE` representation. **B** The sce object is passed to the *iSEE* package using the `iSEE()` function (green box). This function produces an interactive data visualization accessible via a web browser, encapsulating various types of plots into a single application (large light gray box). The various components of the sce data container are used differently by different plots (arrows pointing from slots to plots). The feature assay plot (left) derives its data from the `assays` slot (blue box), whereas the reduced dimension plot (right) derives its coordinates from the `reducedDims` slot (purple box). Clustering information from the `colData` slot is used for each plot to separate and color the data by clusters (middle), as well as to visualize cell metadata (orange box). Cells from any individual plot can be selected via a shiny brush or lasso selection tool, and then transmitted and visualized in any other plot (yellow selection in reduced dimension plot, purple dots on feature assay and column data plot).

#### Interactive Data Visualization

The maturation of web technologies has opened new avenues for the interactive exploration of data. These web technologies have been embraced by the broader R community through the use of *shiny* [147], an R package that facilitates the development of complex interfaces with relative ease. Given the complexity of scRNA-seq data, it is crucial to facilitate the exploration of data across teams of researchers directly working with the data and also to facilitate the communication of results to the broader community. Furthermore, such communication must accommodate audiences with varying levels of programming expertise. To this end, the *iSEE* [148] package provides a full-featured application for the interactive visualization of scRNA-seq data sets through an internet browser, eliminating the need for any programming experience if the instance is hosted on the web (Figure 4). The *iSEE* package directly interfaces with the *SingleCellExperiment* data container to enable the simultaneous exploration of single-cell data and metadata. For example, the *iSEE* package includes functionality to visualize dimensionality reduction results such as *t*-SNE or UMAP while coloring cells, or alternatively, the expression level of any feature by their assigned cluster or cell type. Lastly, the *iSEE* package lets the user export the code required to reproduce the interactive graphics as well, facilitating long-term reproducibility of exploratory analyses.

#### Report Generation

Ensuring that results are well-documented, shareable, and reproducible is of utmost importance for publication and external validation of results. To address this, Bioconductor promotes the usage of R packages like *rmarkdown* [149] and *bookdown* [150] to produce analytical reports, tools which were all used to generate our companion online book (see **Box 2**). Furthermore, Bioconductor has published the *BiocStyle* package [151] to provide standardized formats for package associated vignettes that illustrate software functionality.

Code and reports inherent to a project, especially those associated with documentation or publication, are encouraged to be shared as part of a package as an included vignette as well as in public code repositories such as GitHub. For example, Bioconductor publishes data packages with vignettes that demonstrate how to reproduce an associated manuscript’s relevant figures [152]. In addition, packages such as *iSEE* [148] report all the code used to generate a visualization that was manually specified via the interface described prior.

Beyond bespoke reports or packages, Bioconductor has also added packages that automate the creation of shareable, standalone reports. In particular, one area of focus is the automation of quality control documentation. Namely, the *countsimQC* [153] and *batchQC* [154] packages automate the visualization of various sample characteristics, such as library sizes, the number of genes quantified, and additionally illustrate what batch correction procedures may need to be applied prior to downstream analysis.

### Published and Simulated Data

Access to data – both simulated and published – is essential to validate and benchmark previously established and new methods. Given the rapid ascent of high-throughput single-cell assays, Bioconductor has actively encouraged the community to publish data packages, as well as use standardized data simulation frameworks to foster further development in methods and synthesis of analytic results.

#### Single-cell Data Packages

As new single-cell assays, statistical methods and corresponding software continue to be developed, it becomes increasingly important to facilitate the publication of datasets, both to reproduce existing analysis as well as to enable comparisons across new and existing tools. Bioconductor encourages the publication of data packages, which are primarily focused on providing accessible, well annotated, clean versions of data that are ready to be analyzed by end users.

To standardize the querying of published data packages on Bioconductor, the *ExperimentHub* [155] Bioconductor package was created to enable programmatic access of published datasets using a standardized interface. Amongst available scRNA-seq datasets, the *TENxPBMCData*, and *TENxBrainData* data packages from the 10X Genomics platforms provide streamlined access to the *ExperimentHub* resource for their respective data sets. In addition, the *HCAData* and *HCABrowser* packages provide access to the data portal from the Human Cell Atlas [16] consortium.

It is important to note that with *ExperimentHub* [155], data is processed and submitted by the authors of the package, and thus data packages from multiple sources may not be readily comparable to each other. To address this, single-cell data compendiums, such as *conquer* [112], provide users an interactive online resource of many scRNA-seq datasets where each has been processed in a consistent manner and with additional quality control steps applied. Such an approach helps standardize various pipelines used to process scRNA-seq data in order to make them more readily comparable [112].

#### Benchmarking Datasets

Of note within single-cell data packages, benchmarking datasets are labeled with some known “ground truth” designed for the evaluation of new methods. Amongst these, the *CellBench* [80] data set is one such example specific to scRNA-seq applications, comprised of data from various platforms and cell lines. As such, it can be used to validate batch correction, differential expression, and clustering methods. Another benchmarking dataset for scRNA-seq data is from Tung *et al.* (2017) [156], which can be used for benchmarking methods using data derived from plate-based protocols. Finally, the *DuoClustering2018* package [109] contains data from a variety of sources which were used to test the performance of various clustering methods, with the data primarily chosen to represent different degrees of difficulty in the clustering task.

#### Simulating Data

Simulated data is logical prerequisite for method development where it is necessary to fully know the generative model underlying the data, and thus, accurately benchmark the performance of a method. For scRNA-seq data, the *splatter* package [157] can simulate the presence of multiple cell types, batch effects, varying levels of dropout events, differential gene expression, as well as trajectories, providing a rich canvas for methods testing and development. Furthermore, the *splatter* package uses both its own simulation framework and wraps other simulation frameworks with differing generative models such as *scDD* and *BASiCS* to provide a comprehensive resource for single-cell data simulation.

## Discussion

The open-source and open-development Bioconductor community has developed state-of-the-art computational methods, standardized data infrastructure, and interactive data visualization tools available as software packages for the analysis of data derived from cutting-edge single-cell assays. The rapid development of high-dimensional single-cell assays producing data sets of increasing size, complexity and sparsity, has the Bioconductor community to implement profound changes in how users access, store, and analyze data, including: (1) memory-efficient data import and representation, (2) common data containers for storing data from single-cell assays for interoperability between packages, (3) fast and robust methods for transforming raw single-cell data into processed data ready for downstream analyses, (4) interactive data visualization, and (5) downstream analyses, annotation and biological interpretation. In addition, emerging single-cell technologies in epigenomics, T-cell and B-cell repertoires, and multiparameter assays from single cells (such as joint/simultaneous protein and transcriptional profiling) promise to continue to push forward advances in computational biology. Based on the unique strengths of Bioconductor, in particular its strong connection between users and developers, the high degree of responsiveness of developers to user needs that arise, and the reach of the Bioconductor community, we are optimistic that it will be the hub of development for these technologies as well.

Being a part of the broader R community presents unique advantages for Bioconductor, including accessiblility to statisticians and data scientists. The Bioconductor software project has established best practices for coordinated package versioning and code review. Alongside community-contributed packages, a team of core developers (https://www.bioconductor.org/about/core-team/) implements and maintains the infrastructure needed by the global project, as well as reviews contributed packages to ensure they satisfy Bioconductor package guidelines. Taken together, these practices result in high-quality and consistently maintained packages.

**Table 1:**
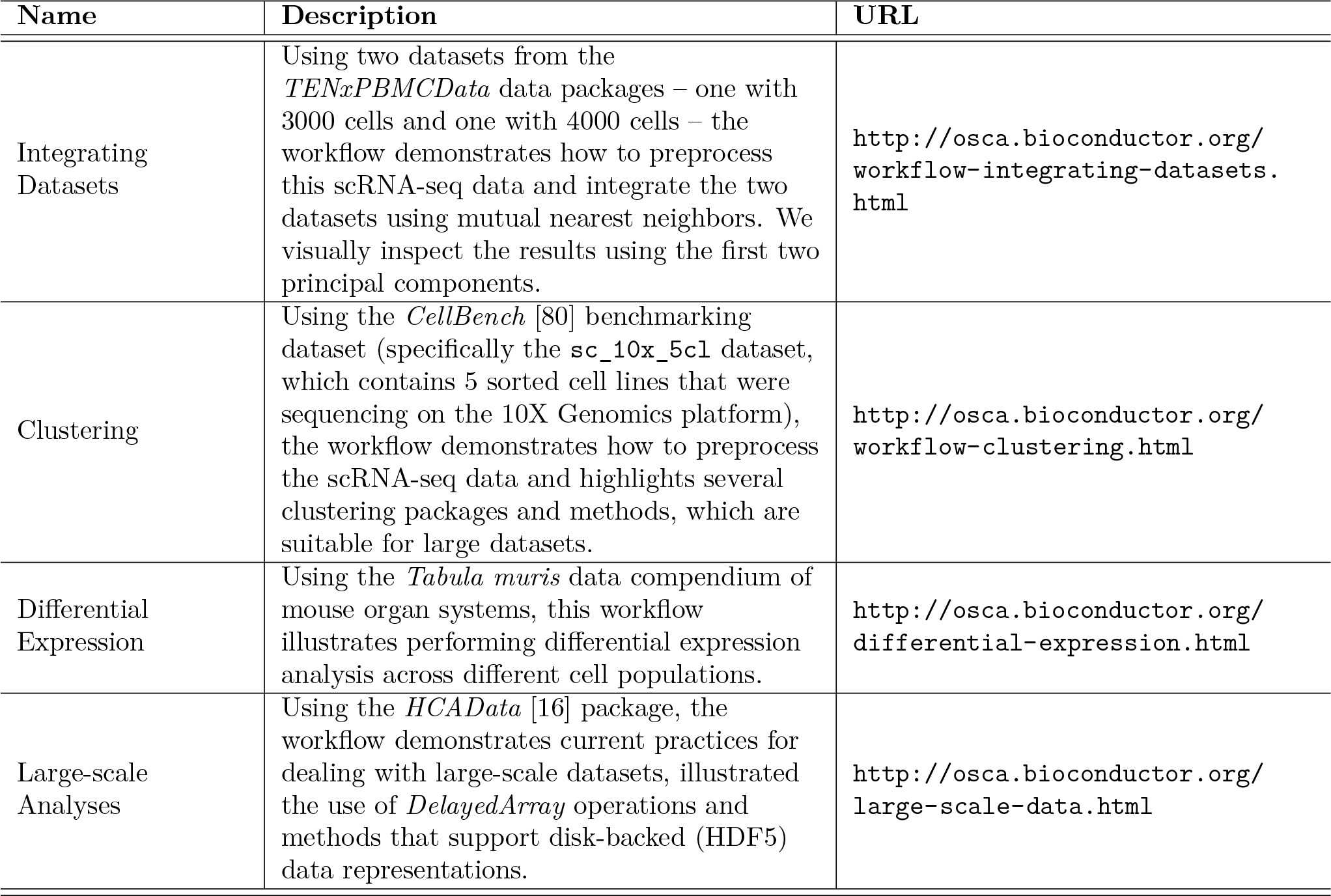
Bioconductor workflows for single-cell analyses.

In addition, Bioconductor provides standardized data containers that enable interoperability between Bioconductor packages, between Bioconductor and CRAN [2] packages, and between R and other programming languages. For instance, it is simple to convert between a *SingleCellExperiment* object and the format used by the popular single-cell CRAN package *Seurat* [158] and Python [159] package *scanpy* [160] and vice versa. Indeed, R has a long history of interoperability with other programming languages. Two notable examples are the *Rcpp* [161–163] package for integrating C++ compiled code into R and the *reticulate* [164] package for interfacing with Python. This interoperability enables common machine learning frameworks such as TensorFlow/Keras to be used directly in R.

To the newcomer, the wealth of single-cell analyses possible in Bioconductor can be daunting. To address the rapid growth of contributed packages within the single-cell analysis space, we have summarized and highlighted state-of-the-art methods and software packages and organized the packages into the broad sections of a typical single-cell analysis workflow (Figure 2) alongside companion code-based workflows showcasing their use (https://osca.bioconductor.org). Finally, Bioconductor software packages are organized into BiocViews, an ontology of topics that classify packages by task or technology. This effort increases discoverability and interoperability. In the future, additional meta packages may be developed to wrap similar methods together.

## Author Contributions

SCH and RG conceptualized the manuscript. RAA, SCH, RG wrote the manuscript with contributions and input from the rest of the authors. All authors read and approved the final manuscript.

## Acknowledgements

Bioconductor is supported by the National Human Genome Research Institute (NHGRI) and National Cancer Institute (NCI) of the National Institutes of Health (NIH) (U41HG004059, U24CA180996), the European Union (EU) H2020 Personalizing Health and Care Program Action (contract number 633974), and the SOUND Consortium. In addition, MM, SCH, RG, WH, ATLL, and DR are supported by the Chan Zuckerberg Initiative DAF (2018-183201, 2018-183560), an advised fund of Silicon Valley Community Foundation. SCH is supported by the NIH/NHGRI (R00HG009007). RAA and RG are supported by the Integrated Immunotherapy Research Center at Fred Hutch. MM is supported by the NCI/NHGRI (U24CA232979). LG is supported by a research fellowship from the German Research Foundation (GE3023/1-1). LW and VJC are supported by the NCI (U24CA18099). VJC is additionally supported by NCI U01 CA214846 and Chan Zuckerberg Initiative DAF (2018-183436). ATLL received support from CRUK (A17179) and the Wellcome Trust (WT/108437/Z/15). FM is supported by the German Federal Ministry of Education and Research (BMBF 01EO1003). MS is supported by the German Network for Bioinformatics Infrastructure (031A537B). DR is supported by the Programma per Giovani Ricercatori Rita Levi Montalcini from the Italian Ministry of Education, University, and Research. HP is supported by the NIH Bioconductor grant (U41HG004059).

## Competing Financial Interests

The authors declare no competing financial interests.

## Supplementary Tables

**Table S1:**
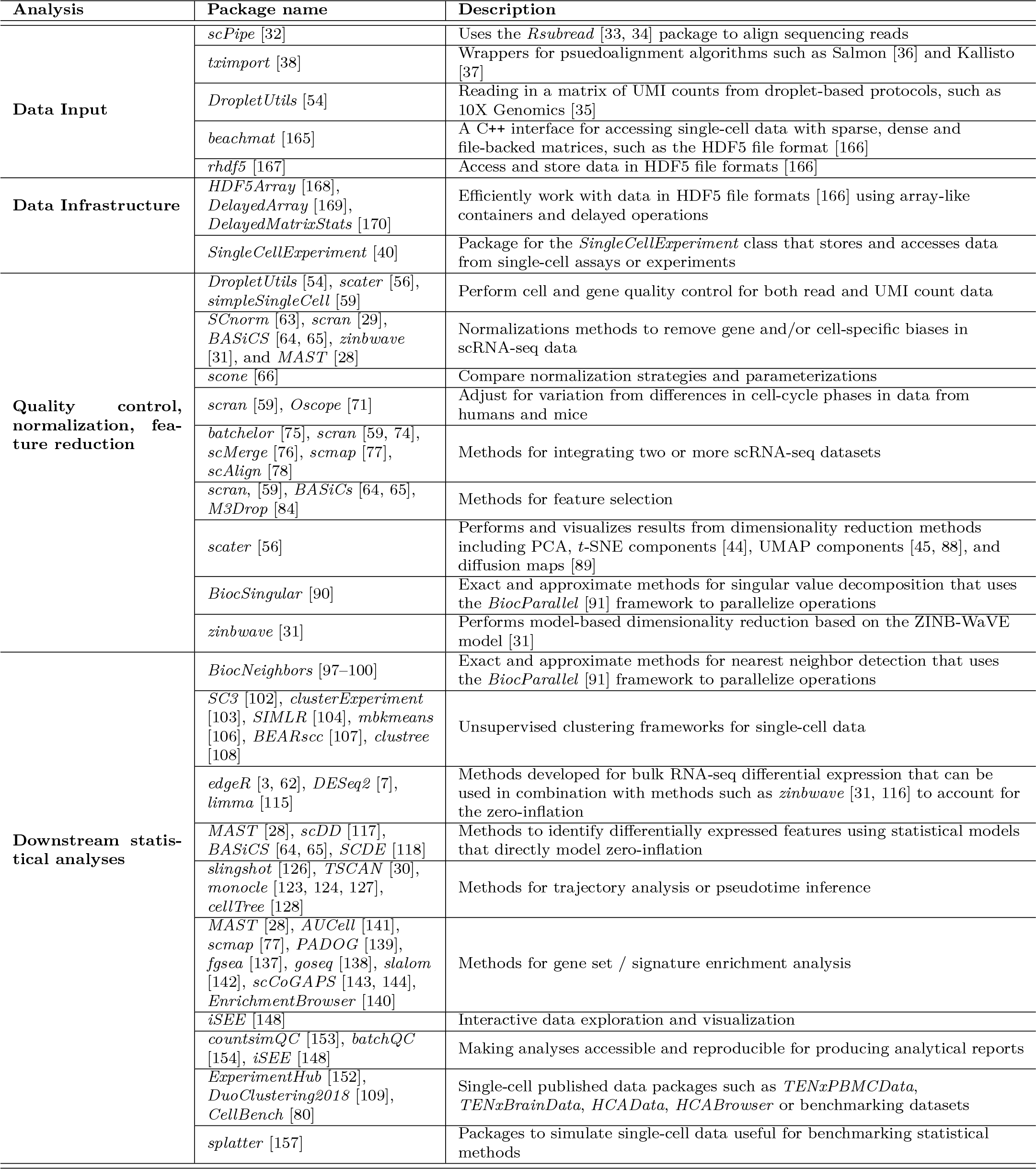
Bioconductor software package for single-cell analyses.

